# Many transcription factor families have evolutionarily conserved binding motifs in plants

**DOI:** 10.1101/2024.10.31.621407

**Authors:** Sanja Zenker, Donat Wulf, Anja Meierhenrich, Prisca Viehöver, Sarah Becker, Marion Eisenhut, Ralf Stracke, Bernd Weisshaar, Andrea Bräutigam

## Abstract

Transcription factors control gene expression during development and in response to a broad range of internal and external stimuli. They regulate promoter activity by directly binding *cis*- regulatory elements in DNA. The angiosperm *Arabidopsis thaliana* contains more than 1,500 annotated transcription factors, each containing a DNA-binding domain that is used to define transcription factor families.

Analyzing binding motifs of 686 and the binding sites of 335 *A. thaliana* transcription factors as well as motifs of 92 transcription factors from other plants, we identified a constrained vocabulary of 74 conserved motifs spanning 50 families in plants. Among 21 transcription factor families, we found one core motif for all analyzed members and between 2 and 72% overlapping binding sites. Five families show conservation of the motif along phylogenetic clades. Five families including the C2H2 zinc finger family show high diversity among motifs in plants, suggesting potential for neofunctionalization of duplicated transcription factors based on the motif recognized. For conserved motifs we tested if they remained conserved since at least 450 million years ago by determining the binding motifs of 17 orthologous transcription factors from 11 families in *M. polymorpha* using amplified DNA affinity purification sequencing. We detected nearly identical binding motifs as predicted from the angiosperm data.

Taken together, the results show a large repertoire of overlapping binding sites within a TF family and species and a high degree of binding motif conservation for at least 450 million years. The results indicate more potential for evolution in *cis-* rather than *trans*-regulatory elements.

## Introduction

Plant transcriptional regulation is a highly diversified process with, for example, around 27,000 nuclear target genes and more than 1,500 transcription factors (TFs) in *Arabidopsis thaliana* (Riechmann et al., 2000). The plant kingdom is highly diverse with about 374,000 existing species (Christenhusz and Byng, 2016), which evolved from an ancestral charophycean alga (Bowman, 2022). Species diversity drastically expanded in the angiosperm lineages 125-112 million years ago (De Bodt et al., 2005) after the colonization of land by streptophytes. In all plants, development and responses to biotic and abiotic challenges require acclimation via changes of gene activity including regulation of transcription. Transcriptional regulation is mediated by sequence-specific TFs, which directly bind DNA. Binding sites (*cis*-regulatory elements) can be characterized *in vivo* and *in vitro*. For *in vivo* characterization, chromatin immunoprecipitation sequencing (ChIP-Seq) is most frequently used, which is influenced by additional *in vivo* factors, such as chromatin structure and partner proteins (Gordân et al., 2009; Li et al., 2011). In contrast, *in vitro* methods, such as protein binding microarrays, high-throughput *in vitro* selection, and DNA affinity purification sequencing (DAP-Seq) allow identification of pure DNA-binding sites without chromatin and methylation influence, but also lack potential binding partners (Berger et al., 2006; Jolma et al., 2013; O’Malley et al., 2016). DAP-Seq also enables comparison of bound sites. Identified binding sites of a TF can be aligned and summarized in a position weight matrix, resulting in a descriptive transcription factor binding motif (TFBM). A TFBM could be for example the G-box CACGTG.

The type and position of binding sites on the DNA in regulatory regions spell out a code, which is read out by the TFs (Seeman et al., 1976; Rohs et al., 2009; O’Malley et al., 2016). Thus, the TFBMs are hypothesized to be a main component to guide the regulatory activity of TFs (Weirauch et al., 2014), but other factors like DNA shape or protein-protein interactions modulate binding sites as well (Appelhagen et al., 2011; Sielemann et al., 2021). Some studies have demonstrated that structurally similar TFs have similar TFBMs (Rushton et al., 1995; Berger et al., 2008; Weirauch et al., 2014; O’Malley et al., 2016; Galli et al., 2018; Lambert et al., 2019) while other studies show that small changes in TF amino acid sequence lead to changes in binding sites and therefore TFBMs (Cook et al., 1994; Noyes et al., 2008; Aggarwal et al., 2010).

TFs are grouped into families based on their shared DNA-binding domains (DBDs) and additional domains, for example for protein-protein interactions (Wilhelmsson et al., 2017). The majority of angiosperm TF families can already be found in the bryophyte *Marchantia polymorpha*, which has not undergone detectable whole genome duplications (Bowman et al., 2017). The divergence of bryophytes and angiosperms can be dated to around 458-467 million years ago (Bowman, 2022), which we will refer to for simplicity as at least 450 million years ago from here on.

TF families in *M. polymorpha* generally consist of fewer members compared to the angiosperm model organism *A. thaliana*. A subsequent increase in TF number is likely due to duplications within families (Catarino et al., 2016). Duplication allows for sequence and function divergence under relaxed selective pressure (Ohno, 1970; Zhang, 2003). Whole genome duplication evidently leads to higher retention rates for duplicated genes by balancing gene dosage effects as opposed to potentially detrimental effects created by single copy duplications (Edger and Pires, 2009; Schmitz et al., 2016). Although most duplicated copies of genes become non-functional within a short evolutionary time by accumulating deleterious mutations leading to pseudogenization (Lynch and Conery, 2000), some gene copies neofunctionalize and adopt completely new functions compared to their paralog. Alternatively, both copies subfunctionalize so that each copy finally covers parts of the function of the original version (Ohno, 1970; Zhang, 2003).

In different origins of multicellularity and therefore in different origins of complex development, different families of TFs expanded and still expand, but the evolutionary source of the TF family frequently predates the expansion event (De Mendoza et al., 2013). Expansion correlates with phylogeny, as phylogenetic branches share specific family expansion patterns (Lang et al., 2010; De Mendoza et al., 2013). In plants, there are several well described examples of TF families that have expanded. The MYB superfamily, for example, can be found across all eukarya, but has increased by 9-fold in member number from the green alga *Chlamydomonas reinhardtii* to *A. thaliana* (Feller et al., 2011; De Mendoza et al., 2013). MYB TFs are defined by their DNA-binding MYB-domain, which consists of a variable number of (imperfect) MYB repeats, each forming three α-helices, with the second and third forming a helix-turn-helix structure to interact with the binding site on DNA. The MYB family can be divided into subfamilies based on the number of MYB repeats (Stracke et al., 2001). MYB-related TFs have a single or partial MYB repeat, which can be either of the R3-type or the R1/R2-type (Dubos et al., 2010). MYB-related TFs are involved in the regulation of diverse functions, including the flavonoid biosynthesis (Dubos et al., 2008) and the circadian clock (Lu et al., 2009).

Very large-scale cross-kingdom analyses have suggested that TF binding specificity to particular TFBMs is predicted by DBD amino acid sequence similarity and that extensive similarity in binding can be detected (Weirauch et al., 2014; Lambert et al., 2019). In *Saccharomyces cerevisiae*, however, 60% of TFs have evolved differential preferences to binding sites due to variations mainly outside of the DBD (Gera et al., 2022). For example, in the zinc finger family in yeast, predominantly intrinsically disordered regions dictate the binding specificity of a TF (Brodsky et al., 2020). In contrast, analyses of TFBMs in *Drosophila melanogaster* and *Homo sapiens* have shown striking conservation between structurally similar TFs despite expansion and divergence of families over the span of 600 million years (Nitta et al., 2015). For example, Activating Transcription Factor-2 (ATF2) has evolved different functions from its orthologue ATF7 in *D. melanogaster* and divergent binding preferences in human and fly were detected (Sano et al., 2005; Nitta et al., 2015). Overall, in animals, extensive conservation of TFBMs is detected in some TF families but not all. The TF family with C2H2 zinc finger as DBD, for example, shows much higher variation in TFBMs despite high protein sequence conservation in metazoans and plants (Lambert et al., 2019). In contrast to animals, most plant genera are tolerant to autopolyploidy and allopolyploidy, which may provide an opportunity for different TF family expansions and for extensive neofunctionalization.

Here, we explored *in vitro* TF binding data from *A. thaliana* TFs and showed that experimentally verified binding of multiple TFs to the same binding site is detectable. We show that TFBMs can be completely conserved within a TF family or be conserved along phylogenetic boundaries within a TF family and quantify the overlapping binding sites. We hypothesized that common TFBMs are a product of TF family expansion and that this conservation dates back to the last common ancestor of angiosperms and bryophytes. To test this hypothesis, we determined the TFBMs of 17 TFs from the bryophyte *M. polymorpha* using ampDAP-Seq.

## Results

To study the experimentally generated binding data of TFs in the genome of *A. thaliana*, we mapped (amp)DAP-Seq peaks from O’Malley et al., (2016) and López-Vidriero et al., (2021) onto individual promoters of genes, here defined as the 1 kb region upstream of the transcriptional start site (TSS) (Figure 1A). Visualizations were amended with accessible chromatin regions from five publications (Lu et al., 2017; Maher et al., 2018; Sijacic et al., 2018; Lu et al., 2019; Sullivan et al., 2019) to visualize accessibility. It is immediately apparent that peak stacks can be observed over defined promoter regions (Figure 1A) showing multiple TFs binding to the same binding site in the promoter. Many of these binding sites are indeed accessible in leaves, root, and whole seedling (Figure 1A). Analyzing 27,206 promoters of nuclear protein coding genes in *A. thaliana*, there is experimental evidence for 0 to 357 (one outlier with 1,076) binding events with a median number of 37 binding events and on average 39 binding events per promoter (-1 kb to TSS; Figure 1B). These numbers represent a lower bound as promoter definitions with increased size increase detectable binding events (Figure 1B). The large positional overlap in TF binding events, which was also qualitatively observed in other promoter regions, suggests large overlap in TFBMs.

**Figure 1.**
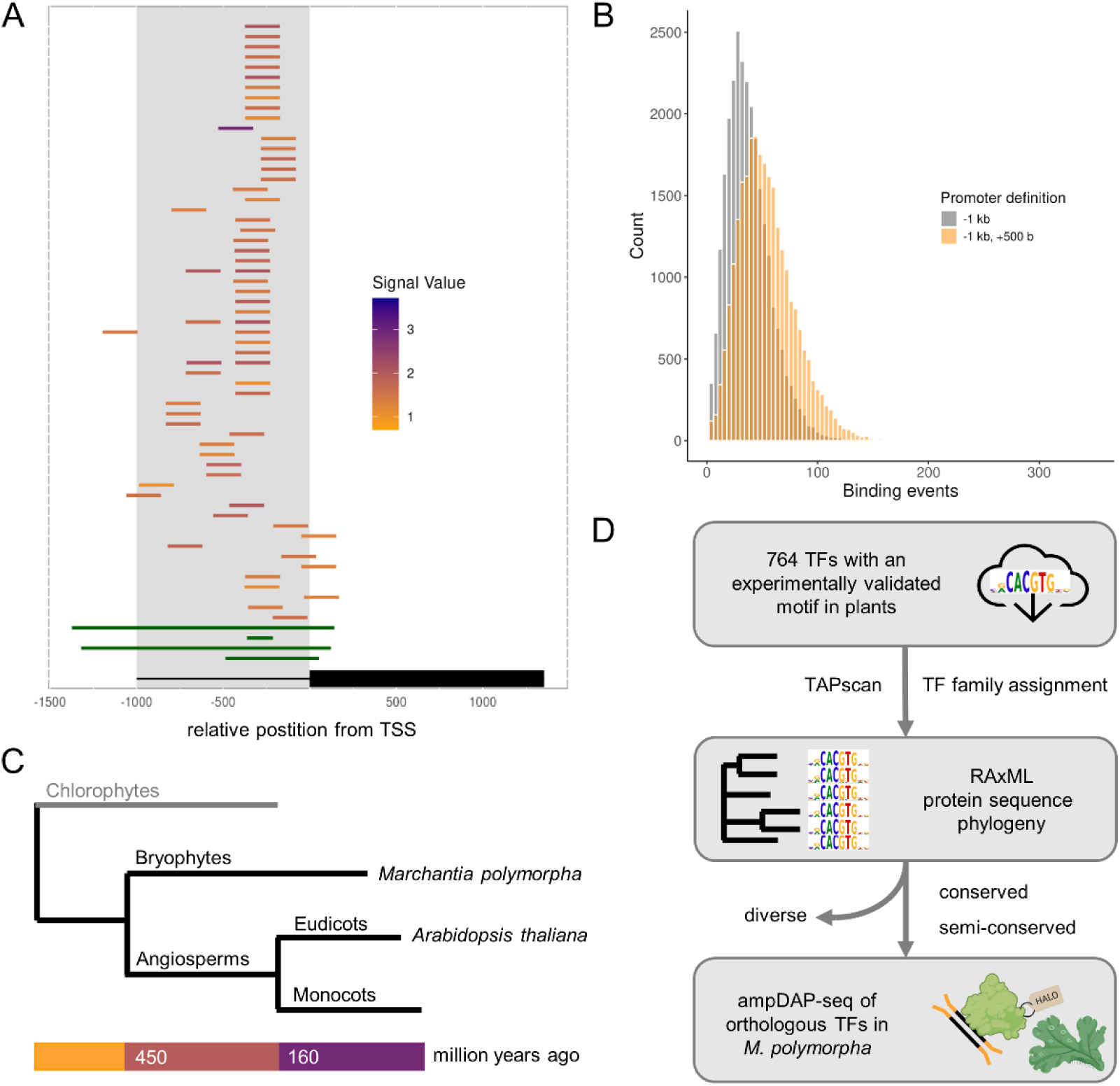
TFBM conservation analysis. A: Representative visualization of (amp)DAP-Seq binding events (peaks), colored from orange to purple by log10-tranformed peak height and open chromatin regions (green) on the *AT2G46310 (CRF5)* promoter. **B:** Histogram showing the number of experimentally determined TF binding sites on two different definitions of nuclear protein coding gene promoters in *A. thaliana* (one outlier each with >1,000 binding sites not shown). **C:** Schematic phylogenetic tree depicting major groups with TFBM data available (black) and chlorophytes as an outgroup (grey). The angiosperm *A. thaliana* and bryophyte *Marchantia polymorpha* are highlighted as representative model organisms. Approximate million years from the last common ancestor below. **D:** Workflow for TFBM conservation analysis. Parts of this figure were created with BioRender.com.

To analyze TFBM similarity within and across TF families in plants, we retrieved all available plant TFBMs in databases (O’Malley et al., 2016; Jin et al., 2017; Castro-Mondragon et al., 2022). To reduce bias, we recalculated the (amp)DAP-Seq derived TFBMs for each TF through a common pipeline, and selected one representative TFBM (Supplemental Table S1) preferring ampDAP-Seq based TFBMs due to the coverage of all genomic binding sites without methylation influence. All TFs with a TFBM were grouped into families based on their DBDs according to TAPscan TF family definitions for *A. thaliana* (Wilhelmsson et al., 2017), and they were aligned and phylogenetically clustered (Figure 1D). We then evaluated each TF family based on how many different TFBMs exist within this family and classified the families into either conserved, semi-conserved, or diverse (see methods for details, Supplemental Table S2).

Database queries yielded a total of 2,190 redundant entries from the Plant Cistrome (O’Malley et al., 2016), JASPAR (Castro-Mondragon et al., 2022), PlantTFDB (Jin et al., 2017) databases. This dataset was amended with DAP-Seq data from López-Vidriero et al. (2021) (Table 1). After removal of redundant and inferred entries, 764 different TFs from 13 different plant species and 50 TF families with TFBM data remained. Data density for plants other than *A. thaliana* is low. Of the 1,725 DNA-binding TFs in *A. thaliana*, 686 have TFBM data available (Table 1). These 686 TFs represent 50 of 71 annotated families in the TAPscan database for *A. thaliana* (Wilhelmsson et al., 2017). Within these 50 families, we know on average 40.5% of TFBMs of all annotated TFs in *A. thaliana*. We tested for overall TFBM similarity and found that the currently known plants TFBMs represent 74 different core TFBMs (Supplemental Table S3), which are between 5 and 21 bp in length with an average length of 8.9 bp (Table 1). Some TFBMs like the WRKY W-box TTGAC are limited to one specific family, while the E-/G-box (C)ACGTG can be found as a TFBM for BES1, bHLH, bZIP TF family members, and one Trihelix family member (Supplemental Table S3).

**Table 1.** Overview of general TFBM statistics from public databases.

|  |  |
| --- | --- |
| Total TFBMs JASPAR | 757 |
| Total TFBMs Plant Cistrome DB | 814 |
| Total TFBMs PlantTFDB | 619 |
| Total TFs (*A. thaliana*) | 1,725 |
| TFs with TFBM (*A. thaliana*) | 686 |
| TFs with TFBM (other species) | 78 |
| Distinct consensus TFBMs | 74 |
| Average TFBM length [bp] | 8.9 |
| TF families (*A. thaliana*) | 71 |
| TF families with TFBM (*A. thaliana*) | 50 |
| TFBMs not assigned to TAPscan family | 18 |
| Average TFBMs known per family (*A. thaliana*) [%] | 40.5 |

### Many families show high levels of TFBM conservation and overlapping binding sites

The TF families with the most TFBMs are the AP2/EREBP family (107) and the MYB family consisting of R2R3- and 3R-type MYB TFs (67). Phylogenetic analyses were performed on complete amino acid sequences for the 32 TF families which contained at least four TFBMs to assess the relationship between amino acid sequence and TFBM similarity. To date the age of a phylogenetic subgroup, we assessed the presence of *M. polymorpha* orthologues determined by Orthofinder2 (Emms and Kelly, 2019) which, if present, indicate an age of at least 450 million years for the branch (Bowman, 2022).

Based on the low number of distinct TFBMs (Table 1) and known conservation for individual families from literature (Ciolkowski et al., 2008; Franco-Zorrilla et al., 2014; O’Malley et al., 2016), we hypothesized that some TF families retain conserved TFBMs despite extensive family expansion. A phylogeny of the WRKY TFs resolved six subclades of which all except clade II-a contain at least one *M. polymorpha* orthologue (Figure 2A). We confirmed the previously described TFBM conservation in the WRKY family (Ciolkowski et al., 2008), with TFBMs available for 45 out of 73 members in *A. thaliana,* all binding the TFBM TTGAC across all clades with minimal variation in the flanking sequences (Figure 2A). Subclade II-d has an additional annotated Zn cluster domain, which does not appear to influence the recognized TFBM (Figure 2A). Based on the conserved TFBM, WRKY TFs could theoretically compete for binding at all TTGAC occurrences and previous analyses have shown substantial overlap of binding sites (O’Malley et al., 2016; Sielemann et al., 2021). To remove potential bias in TFBM detection and to quantify the degree of overlap in actual binding sites, we analyzed the set of all common and distinct binding peaks of 33 WRKY TFs, where (amp)DAP-Seq data is available from O’Malley et al. (2016). Of 35,345 total peaks, 292 are bound by more than 75% of WRKY TFs, 1,568 are bound by one half to 75%, 21,523 are bound by between two and half of all WRKY TFs and 11,962, the minority, is detected as uniquely bound by one WRKY TF (Figure 2C, Supplemental Table S4). To test whether competition can be excluded via different spatial-temporal expression patterns, we compiled expression data from 6,033 *A. thaliana* RNA-Seq experiments and tested for expression similarity by Pearson correlation. Of the 45 WRKY family members with a TFBM in *A. thaliana*, 15 share expression with at least one other family member based on a correlation coefficient of >0.7. Thirty-three have similar expression with at least one other family member based on a correlation coefficient of at least >0.5 (Figure 2D). Contrasting expression patterns indicated by negative Pearson correlation are in the minority (Figure 2D). Examination of the other 31 TF families (Supplemental Data Set, Supplemental Table S2) with at least four characterized TFBMs revealed that 20 additional TF families show extensive TFBM conservation within the TF family (Figure 2B). For example, TFs of the C2C2 GATA zinc finger family unanimously bind the consensus TFBM GATC with little variation in the adjacent bases independent of the method used for TFBM detemination (Supplemental Figure S1). Three orthologues were found in *M. polymorpha*, suggesting potential evolutionarily conservation of this TFBM. Among the conserved TF families, 21 of the 74 different consensus TFBMs (Figure 2B) are represented by 332 TFs.

**Figure 2.**
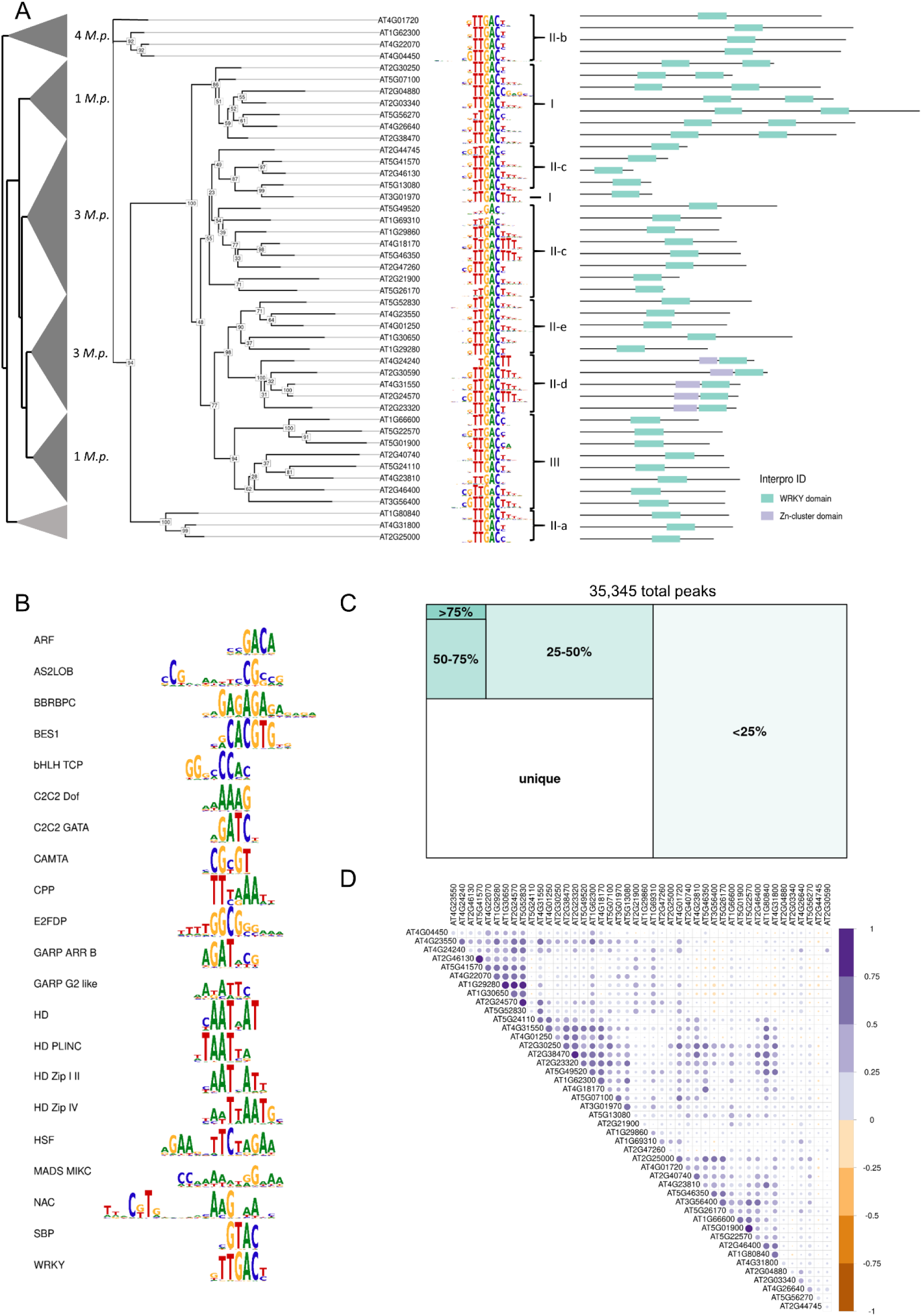
Analysis of TF families with high TFBM conservation. A: Unrooted phylogenetic tree of the WRKY family TFs with TFBMs. Support values at the nodes are based on 1,000 bootstrap iterations. Clade annotations are from Eulgem et al. (2000) and Interpro domain annotations. Collapsed phylogenetic tree is shown with indication of orthologues genes from *M. polymorpha* (*M.p.*) in each subgroup. **B:** Consensus TFBMs of conserved families generated by merging individual TFBMs of TF family members for each TF family. Base height corresponds to information content. **C:** Merged peak set of 33 WRKY (amp)DAP-seq samples showing the subsets of shared and unique peaks by color and subset size. **D**: Expression correlation of 45 WRKY family members in *A. thaliana*. The correlation coefficient is indicated by color and dot size.

A detailed analysis of shared binding sites in each family with a conserved TFBM shows that for all families, sites onto which more than 75% of family members bind, were detected (Supplemental Table S4). For seven families including the C2C2 Dof and the NAC family, the sites with multiple binding TFs also exceed the uniquely bound sites. The other extreme is represented by the BBR/BPC family in which despite the shared TFBM only 253 out of 15,492 detected binding sites are shared between at least two TFs (Supplemental Table S4). The expression divergence of family members within each family, shows that for each of the 21 TFBMs, binding sites and expression patterns overlap to some degree (Supplemental Tables S4, S5). In the analysis, we identified a total of 21 families with one conserved family TFBM (Figure 2B), pointing to a large constraint in the *de novo* evolution of TFBMs as a way for neofunctionalization of TFs in at least these families.

### Semi-conservation of TFBMs follows phylogenetic relationships

Not all TF families are under a similarly strict constraint for the TFBMs of their members. We expected that some families acquired more variation in the TFBM as a potential way to neofunctionalize and regulate different pathways in more than 450 million years of evolution. Our analysis revealed a continuous transition between TF families with completely conserved binding to one single TFBM (Figure 2B) and families in which the TFs bind up to 13 different TFBMs across 33 analyzed members (C2H2 family) (Supplemental Figure S2). Therefore, we established the classification of semi-conservation to cover TF families with up to four different consensus TFBMs and no more than 15% outliers (TFBMs represented by one member). Under this rule, the five families ARID, bHLH, bZIP, MYB, and MYB-related are considered as semi-conserved (Supplemental Table S2).

The MYB-related TF family consists of 62 annotated TFs based on TAPscan for *A. thaliana*, out of which TFBMs for 33 TFs have been experimentally determined, as well as one TFBM each in *Zea mays*, *Solanum lycopersicum,* and *Oryza sativa* (Figure 3A). We observed three different TFBMs in the MYB-related family, GATAA, GATATT, and TAGGG, which generally coincide with phylogenetic subclades of the TF family members. We also observed three outliers in this family, including the determined TFBM from *S. lycopersicum*, which differs from other TFBMs in the phylogenetic tree (Figure 3B). The monocot TFs however recognize the same TFBM as the most similar *A. thaliana* TFs. The TFBMs thus reflect phylogeny of TF family members, highlighting conservation of these TFBMs across the tested plant species. Each of the four major branches in the phylogenetic tree contains at least one *M. polymorpha* TF in the orthogroup indicating that the subgroups have existed since the last common ancestor of bryophytes and angiosperms.

**Figure 3.**
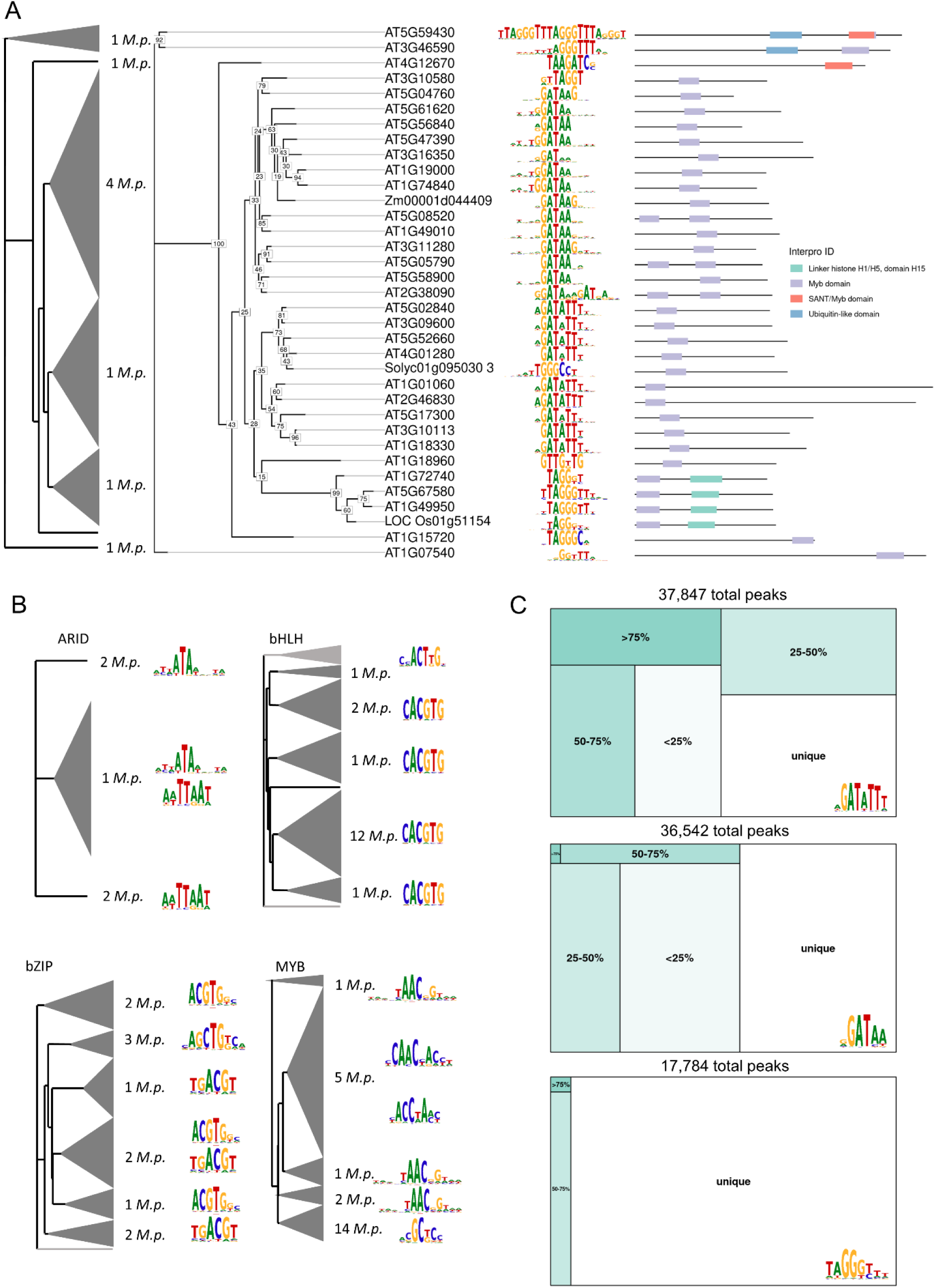
Analysis of TF families with semi-conserved TFBMs. A: Phylogenetic tree of the MYB- related family with support values based on 1,000 bootstraps. Interpro domain annotations indicate structural similarities. Clade annotations are from (Chang et al., 2020) and domain annotations are from Interpro. Collapsed phylogenetic tree with indication of orthologous TFs from *M. polymorpha* (*M.p.*) in each subgroup. **B:** Collapsed phylogenetic relations with orthologues *M. polymorpha* TFs found in the different semi-conserved TF families. Light grey indicates orthologues were not found. TFBMs represent the consensus TFBMs present in the clades. **C:** Merged peak sets for each of the MYB-related TFBM subgroup TF samples showing the subsets of shared and unique peaks by color and subset size.

Within each subclade, the MYB-related TFs potentially compete for binding at binding sites. Expression analysis in *A. thaliana* show that within the 13 members binding the TFBM GATAA, four show similar expression patterns with at least one other family member based on a correlation coefficient of >0.5 (Supplemental Table S5) and 54.7% of all binding sites of the twelve members with DAP-seq data within this subgroup are shared by at least two TFs (Figure 3C, Supplemental Table S4). Among the nine members binding the TFBM GATATT, four share expression patterns with at least one other family member based on a correlation coefficient of at least 0.7 (Supplemental Table S5). The majority of 78.3% of binding sites is not unique (Figure 3C). Among the 7 members binding the TFBM TAGGG, three have similar expression patterns (>0.5) with at least one other family member (Supplemental Table S5). In this subgroup a very large proportion (94.1%) of uniquely bound sites is observed, which are mostly contributed by one of the three TFs (Figure 3C). Although the higher level of divergence of the TFBMs reduces the potential for competition, we detected similar expression patterns between at least two members binding the same TFBM for 40 TFs, indicating that, also in the MYB-related family, different spatial-temporal expression patterns do not exclude the TFs from competition.

The bHLH family members bind two different TFBMs that differ predominantly in two base positions. The initial C of the G-box TFBM CACGTG is missing in the second TFBM and the first G is replaced by a T, leaving the TFBM ACTTG, which was found to be bound by four bHLH TFs (Figure 3B). The bHLH TFs binding to the TFBM ACTTG are more distant from all other members in the tree and the previously analyzed TFs of this subgroup have no Orthofinder2-based orthologues in *M. polymorpha* (Figure 3B). Again, binding site analyses and expression analysis suggests potential competition of different bHLHs for binding at the same binding sites (Supplemental Tables S4, S5). Out of 10,646 binding sites, 4,117 sites are bound by more than one TF. Four of the 28 members binding the CACGTG TFBM subgroup share expression pattern (>0.7) and 13 have similar expression patterns (>0.5). However, the four members with the TFBM ACTTG rarely share binding sites and they have distinct expression patterns with a correlation coefficient <0.5 (Supplemental Tables S4, S5), suggesting binding specificity is potentially established through both site preferences and different spatial-temporal expression.

Grouping of TFBMs along the phylogenetic relationship is also observed in the bZIP family and the MYB family (Figure 3B), suggesting stability of TFBMs during clade-specific expansion. Binding site analyses show that many MYB TF binding peaks and the majority of bZIP binding sites are shared by at least two TFs (Supplemental Table S4). Expression analyses show that there is extensive sharing of expression patterns within members binding the same TFBM (Supplemental Table S5). This group of five semi-conserved TF families covers 18 of the 74 consensus TFBMs including TFBMs that are only bound by one member in a family and were excluded from consensus TFBM generation. The level of competition for each TFBM is reduced compared to the TF families for which the TFBMs are highly conserved with a lower number of TFs who share expression or have a similar expression pattern.

### Diverse TF families bind a variety of different TFBMs

We identified five TF families that have more than four different consensus TFBMs or more than 15% outliers and classified them as diverse. The diverse TF families are ABI3/VP1, AP2/EREBP, C2H2, C3H and Trihelix (Supplemental Table S2). The 188 members in the TF families with less TFBM conservation cover 39 of the 74 identified distinct TFBMs (Table 1). Trihelix TFs have one or two DBDs similar to the MYB domain and are therefore called MYB/SANT-like (Figure 4). The Trihelix family is divided into five clades based on amino acid sequence similarities and the phylogeny largely recapitulates the grouping (Kaplan-Levy et al., 2012) (Figure 4). As of now, TFBMs are available for 14 out of 26 members in *A. thaliana*. All previously defined clades of the Trihelix family have at least one known TFBM (Figure 4). The GT-2 clade has a conserved TFBM of TTTAC. Two TFs in the GT-1 clade, as well as one TF in the SIP1 clade bind to the well-described GT-element with a consensus sequence of GGTTAA (Kaplan-Levy et al., 2012). The third TFBM from the GT-1 clade for TF GT-3a (AT5G01380) is CACGTG (Ayadi et al., 2004), which is also bound by bZIP and bHLH TFs (Figure 3B, Supplemental Figure S3). Comparing transcript level in the leaf and hypocotyl for all 58 TFs with the experimentally determined TFBM CACGTG, we observe similar transcript levels in both tissues (Supplemental Figure S3).

**Figure 4.**
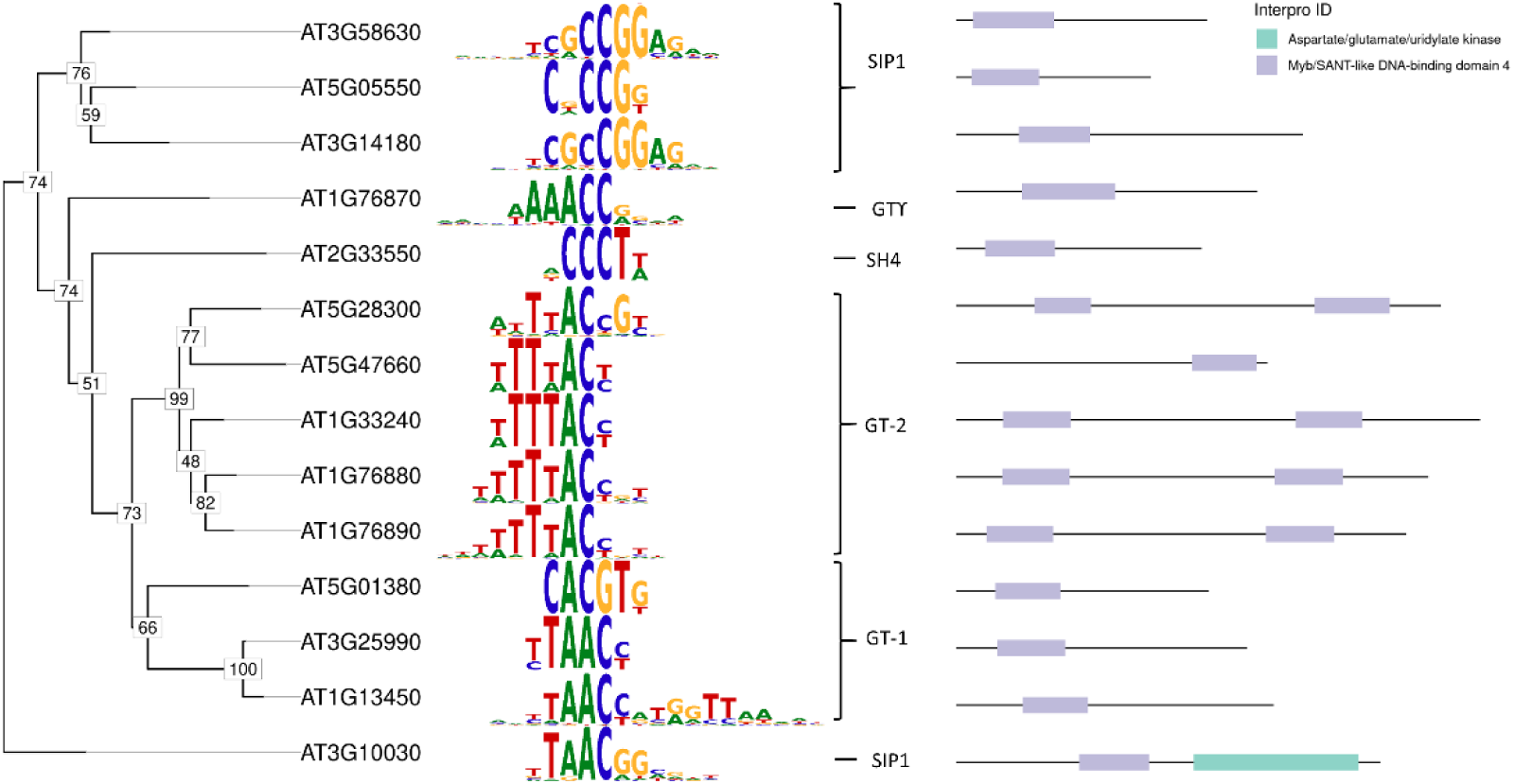
The Trihelix family is an example of a TF family with diverse TFBMs. Unrooted phylogenetic tree of the Trihelix TF family members with TFBM determined. Support values at the nodes are based on 1,000 bootstrap iterations and domain annotations from Interpro.

We also identified a high diversity of TFBMs bound by the C2H2 TF family members in plants (Supplemental Figure S2). Within the C2H2 TF family in plants, a TFBM has been experimentally determined for 32 TFs from *A. thaliana* and one from *S. lycopersicum*. These TFs bind 13 different consensus TFBMs, which is the highest number of different consensus TFBMs within one TF family reported in this analysis (Supplemental Table S2).

### Many TFBMs have been conserved for at least 450 million years

The bioinformatic analyses suggested that TFBMs of particular TF families or subfamilies have been constant since the last common ancestor of bryophytes and angiosperms. To test these hypotheses with laboratory experiments, we selected orthologous TF candidates in *M. polymorpha* as a bryophyte representative and performed ampDAP-Seq to experimentally determine the binding sites *in vitro* and derive the TFBMs.

Of the 14 WRKY TFs in *M. polymorpha* we selected MpWRKY11 and MpWRKY14 for experimental testing. Mapping of bound DNA sequences on the *M. polymorpha* BoGa genome sequence (Beaulieu et al., 2023; https://doi.org/10.4119/unibi/2982437) and peak calling with a negative vector control as background revealed 682 binding sites of MpWRKY11 and 27,902 binding sites of MpWRKY14. The number of peaks loosely correlates with the number of reads per sample, indicating potentially more low affinity binding sites were detected for MpWRKY14. Both experiments yielded a TFBM of TTGAC, which is stable when using only the top 10% of peaks for TFBM detection (Supplemental Figure S6). The NAC family was represented by MpNAC3, which has 41,258 binding sites in the *M. polymorpha* genomic sequences and yielded the gapped TFBM C(G/T)T…AAG, as was predicted from the analyses of *A. thaliana* TFBMs in the TF family. The representatives MpHD3 (HD Zip I-II, 2 members), MpDEL1 (E2F/DP family, 6 members), MpGATA2 (C2C2 GATA family, 3 members), and MpBPCV (BBR/BPC family, 2 members) yielded the TFBMs (A/C)ATNAT, (G/C)GCGGG, GATC and GAGAGA, respectively (Figure 5A). All of these TFBMs were predicted as the likely family TFBM in the analysis of available seed plant TF data (Figure 5A). *M. polymorpha* TFs in the semi-conserved MYB-related family were tested for each TFBM subgroup GATAA, GATATT, and TAGGG. Mp1R-MYB7 was chosen to test for GATAA, MpRVE for GATATT and Mp1R-MYB8 for TAGGG. Mp1R-MYB7 yielded a TFBM of GATNA, MpRVE yielded GATATT, and Mp1R-MYB8 yielded TAGGG with minor changes in the flanking bases (Figure 5B). All TFBMs are highly similar to the TFBMs predicted based on phylogeny (Figure 5B). To pick candidates for testing in the semi- conserved bHLH family, all 42 TFs in *M. polymorpha* were integrated into the phylogenetic tree (Supplemental Figure S4). Although Orthofinder2 did not yield an orthologous TF for the ACTTG TFBM subclade (Figure 3B), bHLH TFs in *M. polymorpha* clustered between *A. thaliana* TFs binding the TFBM ACTTG (Supplemental Figure S5). Therefore, we tested the two *M. polymorpha* TFs MpBHLH27 and MpBHLH44, which yielded C(A/C)ACTTG and CACGTG respectively as predicted based on phylogeny (Supplemental Figure S5). Experimentally determined TFBMs of *M. polymorpha* TFs in the semi-conserved MYB and bZIP family also yielded the TFBM predicted from *A. thaliana* data for one subgroup each (Supplemental Figure S5). Taken together, we detected a highly conserved TFBM within the tested phylogenetic clades for at least 450 million years based on the experimental evidence in *M. polymorpha*. These results support the likely expansion of TFs within the subclades and an evolution of the differential TFBMs predating the split of bryophytes and angiosperms.

**Figure 5.**
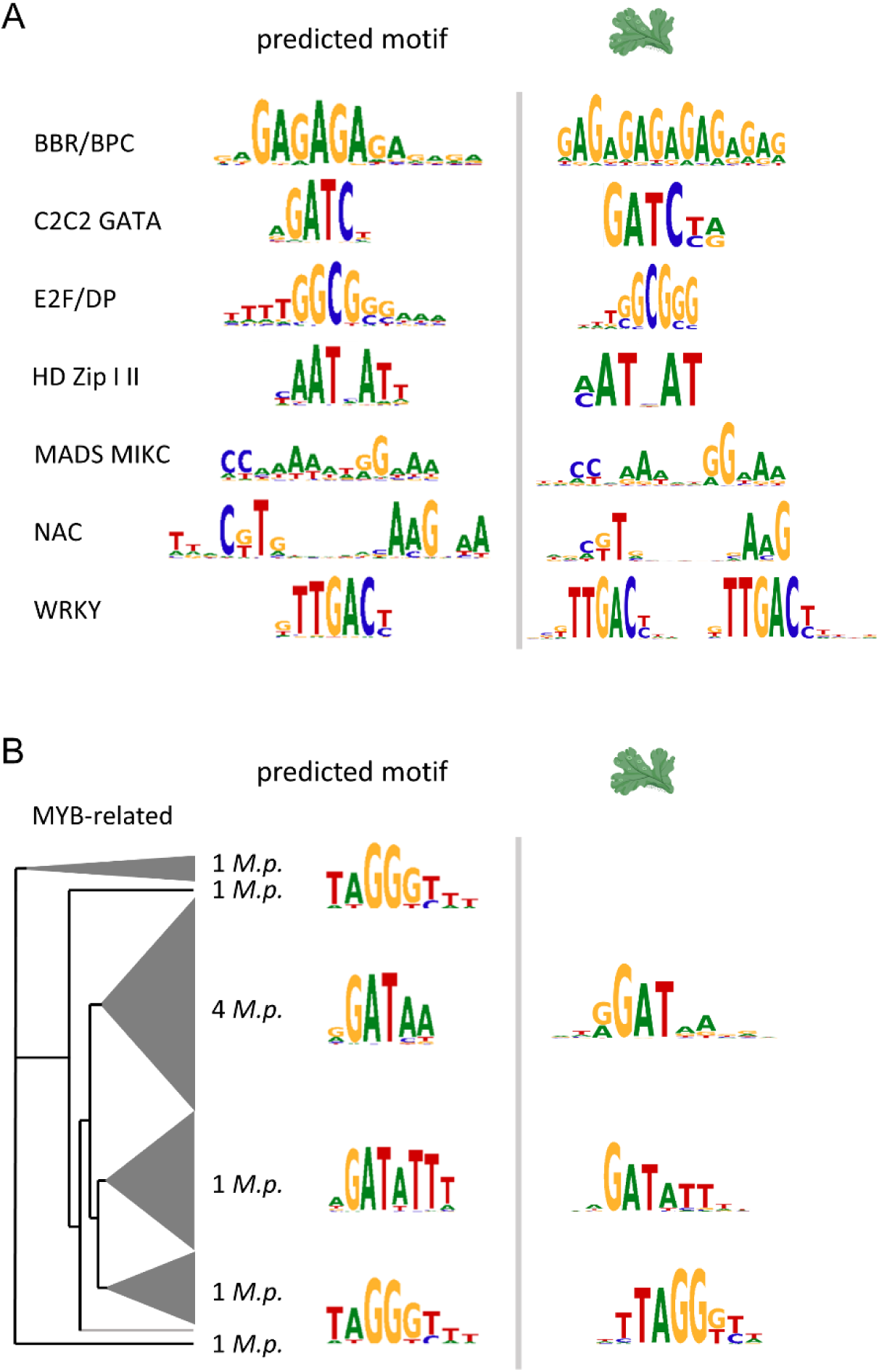
TFBMs in *M. polymorpha* show high conservation in comparison with *A. thaliana* TFBMs. A: Consensus TFBMs determined for each family (left) and experimentally determined TFBMs of one orthologous TF family member in *M. polymorpha* by ampDAP-Seq (right). **B**: Consensus TFBM of each clade (left) and TFBM of one orthologous TF in *M. polymorpha* from each subclade. Parts of this figure were created with BioRender.com.

## Discussion

### There is limited potential for TFBM evolution in some families

DNA-binding TFs of different families have different potential to evolve new TFBMs as a mechanism to neofunctionalize in plants. In families like ARF, BES1, C2C2 GATA, E2F/DP, MADS MIKC, NAC, WRKY and all other conserved families, all member TFs bind the same consensus TFBM (Figure 2B), despite the fact that many of these families have existed at least since the last common ancestor of all land plants. Inclusion of TFBMs directly downloaded from JASPAR and PlantTFDB do not alter the conclusions about conservation indicating TFBMs detection is robust against different technologies (Supplemental Figure S1, also see O’Malley et al. (2016): Figure S1C, Table S1C). This long conservation suggests that, very much unlike the biotechnologically programmable C2H2 TFs (Ichikawa et al., 2023), the defining DBD of these TF families is highly constrained. Conservation may extend to the last eukaryotic common ancestor in some cases as, for example, the strict E2F binding TFBM TTT(G/C)(G/C)CGC in humans (Rabinovich et al., 2008) nearly overlaps with the conserved E2F/DP TFBM TTTGGCG(C/G) determined in *A. thaliana*. The C2C2 GATA TF family, named for (A/C/T)GATA(G/A) as their consensus *cis*-regulatory element sequence in metazoans, binds the TFBM GATC in plants. In *A. thaliana* and *M. polymorpha*, all analyzed members bind to the GATC consensus TFBM, suggesting a shift in TFBMs from the last eukaryotic common ancestor to land plants is possible. GATA TFs contain one or two type IV zinc finger domains and are found in fungi, metazoans, and plants. In fungi and metazoans, C2C2 GATA TFs generally respond to abiotic stimuli (light, nutrients) (Schwechheimer et al., 2022). Inactivating mutations in mammalian C2C2 GATA TFs are associated with developmental diseases and expression of the six TF family members in human is tissue specific to regulate different functions (Tremblay et al., 2018). In plants, a conservation of involvement of C2C2 GATA TFs in control of light-dependent responses is indicated (Reyes et al., 2004). The role of GATA TFs in chloroplast biogenesis in *A. thaliana* was found to not be conserved in *M. polymorpha* (Frangedakis et al., 2024) despite a conserved TFBM (Figure 5A) underscoring evolution in *cis*-regulatory elements rather than in *trans*-acting transcription factors.

On the opposite end of the spectrum, the C2H2 family was reported as diverse in previous studies in metazoans fly and human (Nitta et al., 2015; Lambert et al., 2019), suggesting that some families are inherently more diverse across multiple kingdoms of life. In human, zinc finger clusters need to be able to evolve at a high rate to repress endogenous retroviruses (Lukic et al., 2014). The TF families identified as diverse readily evolve TFBMs *de novo* compared to conserved and semi-conserved families, thus creating possibility for neofunctionalization of TFs via their TFBM.

### Plant specific TF families also show high degrees of TFBM conservation

Some TF families like for example the BBR/BPC, WRKY, NAC, and bHLH TCP family are found in the plant kingdom only (Bowman et al., 2017; Jin et al., 2017; Wilhelmsson et al., 2017). Many semi-conserved TF families (Figure 3) radiated at or before the last common ancestor of all land plants. Thus, it is impossible to determine their degree of conservation by comparison to data from other kingdoms. Within seed plant TF families, we identified a single consensus TFBM in 21 families and semi-conservation along phylogenetic clades in five additional families. Given the presence of *M. polymorpha* orthologues in the subclades of the phylogenetic tree, we hypothesized that the subclades and TFBM differences are at least 450 million years old (e.g., Figures 2, 3).

We tested for conservation in *M. polymorpha* for seven conserved TF families, all subgroups of the semi-conserved MYB-related and bHLH family, as well as one subclade for the bZIP and MYB family each. Unsurprisingly, the E2F/DP family TFBM in *M. polymorpha* matches the TFBM from *A. thaliana* E2F/DP TFs and is a subset of the strict human E2F TFBM. Within the plant specific MADS MIKC, NAC, WRKY, and BBR/BPC TF families, the TFBM determined for *M. polymorpha* TFs matches the family TFBM determined in seed plants (Figure 5A). These plant specific TF families are thus similarly constrained as the E2F/DP TFs (Rabinovich et al., 2008) in their neofunctionalization with regard to their TFBM. This constraint appears true even though the binding site specificity varies greatly with BBR/BPC TFs binding rather specifically in the seed plant *A. thaliana* while the other families show large overlap in actual binding sites between their members (Supplemental Table S4).

The duplication of TFs in families with conserved TFBMs, however, has not enabled evolution of new TFBMs as a mechanism for neofunctionalization (Figure 5A). The analysis of all phylogenetic TFBM subgroups of the semi-conserved MYB-related family shows that each subgroup contains at least one *M. polymorpha* orthologue (Figure 3A) and that the tested orthologue from each subgroup matches or nearly matches the TFBM (Figure 5B). While these TF families are able to evolve new TFBMs, they do so only very rarely. The data suggests that they did not evolve a new TFBM since the last common ancestor of all land plants approximately 450 million years ago despite an increase of TF family members (Figure 5B, Supplemental Figure S4, S5). It is tempting to speculate that the colonization of a new and empty niche despite the many challenges or maybe because of the many challenges enabled radiation among the usually constrained TF families. The presence of what are called outliers here, that is TFBMs, which occur only once within a TF family, such as a TFBM CATCAT among the MADS MIKC TF family (Supplemental Data Set), may either represent technical errors, such as mislabeling or co-occuring binding sites under the peak, or potentially represent a novel TFBM evolution within seed plants.

Since the last common ancestor of all land plants, seed plants have evolved a number of innovations, such as flowers, seeds, pollen, and secondary growth among others. (Bowman et al., 2017). Many of the TFs known to control the trait in the seed plant *A. thaliana* are already present in the last common ancestor of all land plants. For example, the MADS MIKC TF AGAMOUS-LIKE 2 is involved in flowering in *A. thaliana* and its orthogroup member in the non-flowering liverwort *M. polymorpha* MpMADS2 already binds the MADS typical “CArG box” (Figure 5A, Supplemental Data Set). GATA TFs in *A. thaliana* are also involved in flowering (Richter et al., 2013) and have thus acquired novel functions since the last common ancestor of *M. polymorpha* and *A. thaliana* despite no changes in the conserved TFBM GATC (Figure 2B and Figure 5A). The hypothesis of a new “master regulator” for a new trait, such as posited, for example, for C4 photosynthesis (Westhoff and Gowik, 2010) is tempting. However, the extensive TFBM conservation among many TF families (Figures 2, 3, and 5) suggests that, if at all, evolution of a new TFBM is more likely in diverse families (Figure 4). We postulate that the likelihood of changes in *cis*, that is to the promoter syntax of target genes, to enter an existing regulon (van den Bergh et al., 2014) is higher. The observed conservation of TFBMs between bryophytes and angiosperms (450 million years) echoes a similar finding in metazoans (600 million years) (Lambert et al., 2018) and yeast (Nitta et al., 2015). As in animals, we observe a number of diverse TF families *in planta* that have evolved multiple TFBMs over time (Figure 4).

### Multiple factors mediate binding specificity of TFs

While both anecdotal and large-scale systematic analyses of TFBMs suggested that phylogenetically related TFs bind the same or a very similar TFBM (Figure 2A, B) (Weirauch et al., 2014; O’Malley et al., 2016; Lambert et al., 2019; Tu et al., 2020), mutant analysis of TF knockout mutants suggest that phylogenetically related TFs have very different functions. For example, TFs of the R2R3-MYB family are involved in many processes including development, regulation of flavonoid biosynthesis, and resistance despite conserved DBDs (Stracke et al., 2001; Dubos et al., 2010). The similarity of TFBMs makes it difficult to conceptualize specific transcriptional regulation. Although *A. thaliana* contains over 1,700 TFs, transcriptional regulation operates with a limited vocabulary of 74 TFBMs based on 686 TFs analyzed experimentally (Table 1). Similar small numbers of 72 consensus TFBM based on 619 TFs from *A. thaliana* (Jores et al., 2021) and 85 consensus TFBMs based on 529 *A. thaliana* TFs (O’Malley et al., 2016) have been reported previously based on slightly different methods to compare TFBMs. Potentially, DNA shape conferred by TFBM adjacent bases yields specificity despite conserved TFBS (i.e. Rohs et al., 2009; Gordân et al., 2013; Sielemann et al., 2021). Here, we show that a binding site complements the TFBM analysis and demonstrates extensive overlap in binding (Figures 2C and 3C, Supplemental Table S4). While a minority of families and subfamilies, notably the BBR/BPC and the MYB-related TAGGG, have more than 90% uniquely bound sites despite shared TFBMs, the majority of TF families and subfamilies show large overlaps in TFBMs and binding sites. The reported overlap in binding sites provides a lower bound, as deeper sequencing might lead to detection of additional binding sites.

The proportion of overlaps with more than half of the members of a family binding a particular site demonstrates that the overlap is not a function of the most recent whole genome duplication event (Figures 1A, 2C, 3C, Supplemental Table S4) (Bowers et al., 2003).

In principle, specificity may also be achieved via different spatial-temporal expression patterns (Breuninger et al., 2008), different partner proteins (Kim et al., 2008; Appelhagen et al., 2011), histone modifications (Charron et al., 2010; Zhao et al., 2019), chromatin accessibility (Lu et al., 2019), and binding site positioning and spacing (O’Malley et al., 2016; Galli et al., 2018). Here, we explored whether expression patterns between TFs that share a TFBM differ using 6,033 *A. thaliana* wildtype RNA-Seq experiments. In all cases tested, at least one and frequently multiple TFs, which share a TFBM also share their expression patterns to some (correlation >0.5) or to a large degree (correlation >0.7) (Supplemental Table S5) indicating that different spatial-temporal expression only contributes a minor share to specificity in the majority of cases. Chromatin accessibility may also only play a minor role in mediating specificity as TFs not only share a TFBM but also share binding sites (Figures 1A, 2C, and 3C, Supplemental Table S4). The DNA shape contributes to binding ability and to binding specificity (Sielemann et al., 2021) but divergence is not absolute (Sielemann et al., 2021). Binding site spacing and protein-protein interactions were not explored in this work. For a subset of bHLH MYC-related TFs, a relative but not absolute preference for binding site spacings is known (López-Vidriero et al., 2021). If we assume that both effects in general are similarly small compared to expression patterns, DNA shape and chromatin accessibility, the overlap in binding sites and expression patterns (Figures 2 and 3, Supplemental Tables S4 and S5) opens up the possibility that TFs compete for the same binding site.

Competition at binding sites provides an explanation for the high variation in complementation experiments observed for TFs on binding sites, which are bound by many partners (Lee et al., 2007; Stracke et al., 2010; Gangappa and Botto, 2016). Competition at binding sites and different binding affinities may also provide one explanation for the still substantial gap in predictability of gene expression from sequence and from binding events alone (Li et al., 2018; Avsec et al., 2021; de Almeida et al., 2022). It may also explain why TFs counterintuitively act as both activators and repressors (Mahendrawada et al., 2023), since, intriguingly, HY5 experiments with the TF carrying an enhancer or repressor domain still fail to clarify if native HY5 acts as a repressor or activator on its targets (Burko et al., 2020).

### Conclusions

Taken together, our results show that TFs in plants act on a limited vocabulary of TFBMs, which has been conserved for at least 450 million years in selected families and act on overlapping binding sites. In conserved and semi-conserved families, further conservation of TFBMs along phylogenetic clades can be predicted from on phylogenetic analyses of available plant TFBMs and overlapping binding sites detected. The combination of limited neofunctionalization of TFs via variant TFBMs with the evolution of major innovations in seed plants supports the hypothesis that most evolution must occur in *cis-* rather than *trans*- regulatory elements, at least if the innovations are regulated by TF families with conserved or semi-conserved TFBMs. Competition at binding sites may play a larger role than previously appreciated.

## Methods

### Amplified DNA affinity purification sequencing (ampDAP-Seq)

Plants were grown for six weeks on half-strength Gamborg’s medium (Gamborg B5; Duchefa Biochemie B.V., Netherlands) in petri dishes with a 16h/8h light/dark cycle at room temperature. DNA was extracted from male G2 generation *M. polymorpha* subsp. *ruderalis* BoGa (Busch et al., 2019) with the cetyltrimethylammonium bromide (CTAB) method (https://dx.doi.org/10.17504/protocols.io.bcvyiw7w).

DNA (4-5 µg) was fragmented by sonication to 200 bp with the M220 Focused-Ultrasonicator (Covaris, USA). End-repair, A-tailing and Y-adaptor ligation were performed following the protocol of (Bartlett et al., 2017): For the sample clean-up the DNA was purified using AMPure XP beads (Beckman Coulter, USA) instead of ethanol precipitation and the NEBNext® End Repair Module for end-repair. To obtain an ampDAP library, 15 ng of the DAP library was amplified with 11 cylces PCR. For the binding assay, genes were cloned in pFN19A (N-terminal HaloTag®) T7 SP6 Flexi® vector (Promega, USA; discontinued) by Gibson assembly (Gibson et al., 2009). All primer sequences are available in Supplemental Table S6. TFs as well as an empty HaloTag vector control were expressed with TnT® Coupled Wheat Germ Extract System (Promega, USA) using 2 µg plasmid DNA. Halo-fusion proteins were purified with Magne® HaloTag® Beads (Promega, USA) and then incubated with 50 ng ampDAP library. DNA was recovered, amplified and indexed with 16-20 PCR cycles. Fragments between 200 and 400 bp were extracted from a 1% agarose gel with QIAquick Gel Extraction Kit (Qiagen, Netherlands). The final library was sequenced as 85 bp long single-end reads on a NextSeq™ 550 or NextSeq™ 2000 (Illumina, USA).

### TFBM determination

TF families and their members in *A. thaliana* were retrieved from TAPscan (Lang et al., 2010; Wilhelmsson et al., 2017). DAP- and ampDAP-Seq data from O’Malley et al. (2016) was obtained from the Gene Expression Omnibus (GEO) database (Barrett et al., 2013) under the accession GSE60143 and data from López-Vidriero et al. (2021) under GSE155321 and analyzed according to Sielemann et al., (2021): peak sequences were extracted from the TAIR10 reference genome (https://www.arabidopsis.org) of *A. thaliana* and TFBMs were determined using MEME-ChIP (Machanick and Bailey, 2011). The TFBM with the lowest e-value and less than 21 bases was chosen to avoid long artifacts. AmpDAP-Seq data from *M. polymorpha* was mapped to the reference genome *M. polymorpha* subsp. *ruderalis* BoGa (https://doi.org/10.4119/unibi/2982437) using Bowtie v.2.4.2 (Langmead and Salzberg, 2012). Output files were converted to BAM format with SAMtools (Danecek et al., 2021) view and sorted (samtools sort -n). To remove duplicates, the SAMtools commands fixmate, sort and markdup were executed with default parameters. Peaks were called using GEM (Guo et al., 2012) or MACS3 (Zhang et al., 2008) including the empty vector sample as a control. Peak sequences centered around the peak summit or the top 10% of peaks ranked by the fold enrichment (SignalValue) were submitted to MEME-ChIP for TFBM extraction and yielded the same TFBMs (Supplemental Figure S6). Additionally, we used experimentally determined TFBMs directly from the open access databases JASPAR2022 Plant Core (Castro-Mondragon et al., 2022) and PlantTFDB (Jin et al., 2017), if no DAP-Seq data was available. All used TFBMs for analyses in this study with their source and the original experimental method are listed in Supplemental Table S1.

### Family allocation

All TFs were assigned to TAPscan TF families in *A. thaliana*. If a corresponding TF family in another database exists in TAPscan, annotations were converted. Accurate allocation was validated via annotated Interpro domains (Blum et al., 2021). TFBMs from other species were assigned to the TF families using blastp (Altschul et al., 1990) with the corresponding protein sequence against the *A. thaliana* TAIR10 proteome (https://www.arabidopsis.org). The protein sequences of the TFs were retrieved from Uniprot or the respective proteome from Phytozome (Goodstein et al., 2012) if available. The best blast hit ranked by e-value and secondly percentage of identity was chosen to determine the nearest *A. thaliana* orthologue, whose TF family annotation was then transferred. The databases contain 18 TFBMs, which could not be assigned to a distinct TAPscan TF family. The MYB family was manually reduced to contain only MYB3R and R2R3-MYB factors in accordance with annotations from Table 1 in Stracke et al., (2001) and Plant Cistrome database annotations. Some TFs had multiple TFBMs generated by different methods, requiring the selection of one representative TFBM per protein. We preferred ampDAP-Seq data over DAP-Seq data followed by other determination methods, because DAP-Seq is a high throughput method capturing the most complete set of binding sites on gDNA (O’Malley et al., 2016) and ampDAP-binding is solely based on sequence and not influenced by methylation state. We reassigned the JASPAR TFBMs originally retrieved from ReMap to the original method of either ChIP-Seq or DAP-Seq.

### Phylogenetic analyses

Full protein sequences were aligned per family using MUSCLE v3.8.31. Phylogenetic trees were generated with RAxML v7.4.4 with 1,000 bootstraps and the PROTGAMMAJTT matrix (method modified from Guedes Corrêa et al., 2008) for similarity measure. The phylogenetic trees with TFBMs were visualized using the R package motifStack (Ou et al., 2018) (Supplemental Data Set). Protein sequences were scanned for domain annotations from Pfam and Prosite using InterProScan (Jones et al., 2014).

The similarity of TFBMs within a family was assessed using compare_motifs from the R package universalmotif with default parameters. Clusters were generated by cutting a dendrogram resulting from the distances of the TFBMs at 0.5 followed by manual curation (method modified from Jores et al., 2021). Since gapped TFBMs are more difficult to detect (Bailey et al., 2009), we considered sufficiently similar parts of a gapped TFBM as the same cluster. Consensus TFBMs were generated with mergeMotifs (motifStack) and for gapped TFBMs merge_motifs (universalmotif) for each cluster and trimmed with trim_motifs (universalmotif). We defined TFBMs appearing only once in a TF family as outliers and excluded them from consensus TFBM generation (method modified from Jores et al., 2021). Distinct TFBMs across all TF families were established by TFBM comparison as described before for individual families and the clusters generated from an hclust analysis in R and cutting the tree with a cut-off of h=0.2.

We considered TF families with at least four TFs with a known TFBM for our assessment of conservation level within the family, as this is the cutoff for the generation of a phylogenetic tree with RAxML. Families with one consensus TFBM, subtracting outliers, and less than 15% outlier TFBMs were considered as a conserved family. Up to four different consensus TFBMs and less than 15% outliers classified families as semi-conserved (see Supplemental Table S2).

Orthologues proteins in the bryophyte *M. polymorpha* were determined using OrthoFinder2 (Emms and Kelly, 2019) for the TFs with a TFBM in a given phylogenetic clade of the tree.

### Additional analyses

Wild-type RNA-Seq experiments of *A. thaliana* were downloaded from the SRA (Leinonen et al., 2011) and mapped onto the TAIR10 reference genome (as described in Halpape & Wulf et al., 2023). Pearson correlation was calculated within conserved families and within groups binding to the same TFBM in semi-conserved families and visualized using the corrplot R package. Expression levels in the leaf and hypocotyl of *A. thaliana* from Klepikova et al. (2016) were averaged across the two replicates and log2 transformed.

Common and distinct peaks between TFs with the same TFBM within families were identified using Homer (Heinz et al., 2010) mergePeaks -d given. Peak sets were visualized using the treemaps R package.

## Data availability

Raw ampDAP-Seq data for *M. polymorpha* subsp. *ruderalis* BoGa is available under the Bioprojects PRJNA1007631 and PRJNA1148438 on the NCBI SRA.

All code used in this study, as well as all consensus TFBMs and individual TFBMs in MEME-format are available on GitHub (https://gitlab.ub.uni-bielefeld.de/sanja.zenker/tfbm-evolution) (will be made available upon publication).

## Acknowledgements and Funding

This study was funded by the German Research Foundation (Deutsche Forschungsgemeinschaft; DFG) through the Sonderforschungsbereich 175 (SFB-TRR175): “The Green Hub, Central Coordinator of Acclimation in Plants”, and through “Evolutionary network analysis based on the transcriptome atlas of *Marchantia polymorpha*” Funding ID: BR4617/1-1. We gratefully acknowledge support by the BMBF-funded de.NBI Cloud within the German Network for Bioinformatics Infrastructure, and the CeBiTec compute cluster for computational resources.

## Author Contributions

SZ analyzed the data and co-wrote the manuscript, DW conceived of the study, AM and SZ produced and analyzed ampDAP-Seq data in *Marchantia polymorpha*, PV performed ampDAP-Sequencing, SB re-analyzed HY5 experiments, ME edited the manuscript, RS edited the manuscript, BW edited the manuscript, AB suggested analyses and co-wrote the manuscript.

